# Development of a Finite Element Model to Predict the Cellular Micromechanical Environment in Tissue Engineering Scaffolds

**DOI:** 10.1101/2020.06.25.170597

**Authors:** Mitchell I Page, Peter E Linde, Christian M Puttlitz

## Abstract

Cell fate in tissue engineering (TE) strategies is paramount to regenerate healthy, functional organs. The mechanical loads experienced by cells play an important role in cell fate. However, in TE scaffolds with a cell-laden hydrogel matrix, it is prohibitively complex to prescribe and measure this cellular micromechanical environment (CME). Accordingly, this study aimed to develop a finite element (FE) model of a TE scaffold unit cell that can be subsequently implemented to predict the CME and cell fates under prescribed loading. The compressible hyperelastic mechanics of a fibrin hydrogel were characterized by fitting unconfined compression and confined compression experimental data. This material model was implemented in a unit cell FE model of a TE scaffold. The FE mesh and boundary conditions were evaluated with respect to the mechanical response of a region of interest (ROI). A compressible second-order reduced polynomial hyperelastic model gave the best fit to the experimental data (C_10_ = 1.72×10^-4^, C_20_ = 3.83×10^-4^, D_1_ = 3.41, D_2_ = 8.06×10^-2^). A mesh with seed sizes of 40 μm and 60 μm in the ROI and non-ROI regions, respectively, yielded a converged model in 54 minutes. The in-plane boundary conditions demonstrated minimal influence on ROI mechanics for a 2-by-2 unit cell. However, the out-of-plane boundary conditions did exhibit an appreciable influence on ROI mechanics for a two bilayer unit cell. Overall, the developed unit cell model facilitates the modeling of the mechanical state of a cell-laden hydrogel within a TE scaffold under prescribed loading. This model will be utilized to characterize the CME in future studies, and 3D micromechanical criteria may be applied to predict cell fate in these scaffolds.

## 1 Introduction

Tissue engineering (TE) aims to restore the healthy function of organs by stimulating progenitor cells into a regenerative process. To this end, the fate of these cells is critical. In many tissues, the cellular micromechanical environment (CME) largely drives these cell fates (1–4). However, it is prohibitively complex to prescribe and measure the inhomogeneous, three-dimensional CME in the hydrogel matrix of tissue engineering scaffolds. Accordingly, a fundamental understanding of how the loading, architecture, and materials of TE scaffolds influences the CME has not been comprehensively described.

Computational models, such as those that utilize the finite element (FE) method, hold the potential to increase our understanding of mechanobiological responses for a broad range of scaffold and target tissues (5–7). A previously-developed FE model of a TE scaffold has replicated the important mechanical properties of the intervertebral disc’s annulus fibrosus (8), however, the CME generated within this scaffold under physiological loading remains unclear. Accordingly, the aim of this work was to: (1) characterize the mechanical behavior of a common fibrin hydrogel and (2) to implement this hydrogel as the composite matrix material within the previously-developed model. A unit cell model of a repeating lattice architecture was created and the model mesh and boundary conditions were evaluated. Overall, this work created a computational unit cell model that can be utilized to predict the CME in TE scaffolds.

## 2 Materials and Methods

### 2.1 Mechanical characterization of fibrin hydrogel

#### 2.1.1 Fabrication of fibirn samples

Whole blood was collected from healthy, mature sheep and fibrinogen was isolated using the Cohn fractionation method (9). Whole blood collection was performed under approval from the Institutional Animal Care and Use Committee at Colorado State University (protocol #: KP104). Fibrin hydrogel samples were cast into cylindrical molds (15.9 mm diameter) by pipetting 1.2 mL of fibrinogen into the mold, adding 4 mg of thrombin (bovine thrombin, MilliporeSigma, Burlington, MA, USA) and mixing the fibrin and thrombin by gently pipetting in and out of the mold. The fibrin samples were solidified at room temperature, maintained at 37°C for 24 hours, then gently removed from the molds (Figure 1a).

**Figure 1.**
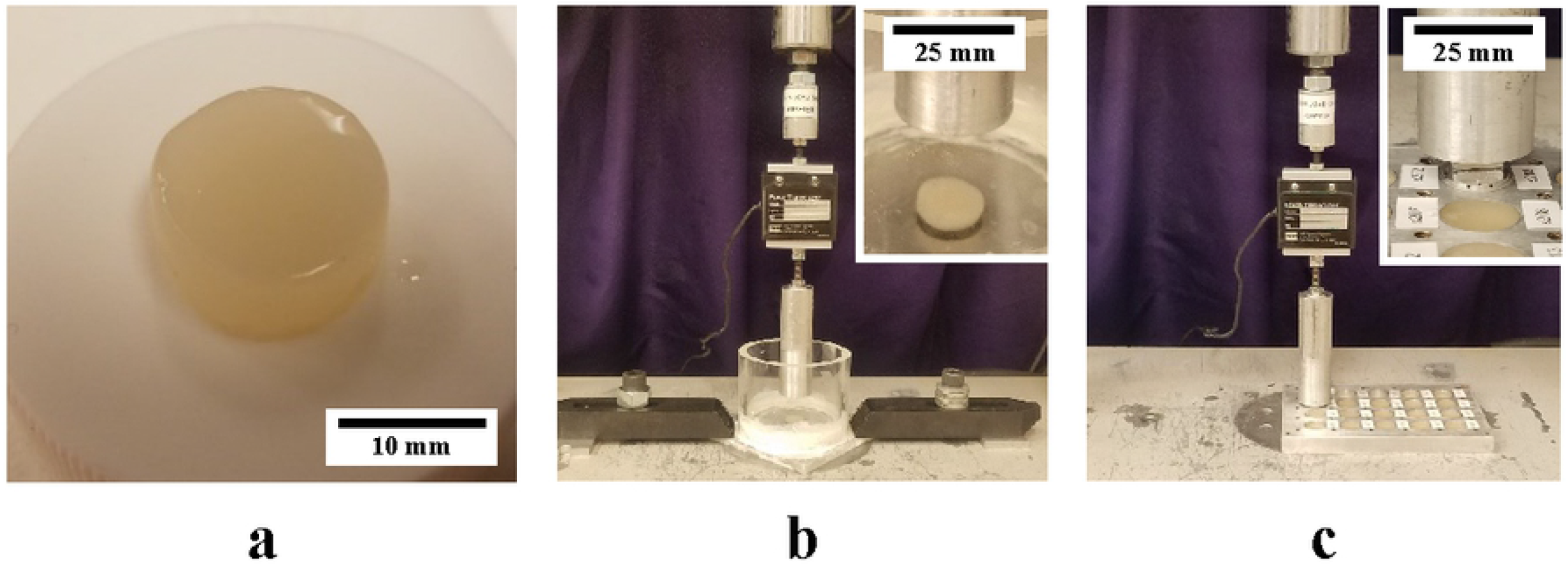
Mechanical testing of fibrin hydrogels: (a) cylindrical sample of fibrin hydrogel removed from mold, (b) experimental apparatus for unconfined compression, and (c) experimental apparatus for confined compression. Insets show close up images of the samples in the experimental configurations.

#### 2.1.2 Material testing of fibrin

Hydrogel specimens were divided into two groups for unconfined compression (n = 9) and confined compression (n = 7) testing. Prior to testing, the diameter and height of each specimen was measured with digital calipers (CD-6”PSX, Mitutoyo, Kawasaki, Japan). Unconfined compression and confined compression tests were conducted using a servo-hydraulic materials testing system (858 MiniBionix; MTS Systems Corp., Eden Prairie, MN) with a 100 N force capacity tranducer (Model 661.09B-21, MTS Systems Corporation, Eden Prairie, MN, USA) and a custom designed testing apparatus (Figure 1b-c). Samples were preconditioned with 10 cycles of 5% strain then compressed to 50% strain at a displacement rate of 0.1 mm/s. Force and displacement data were recorded at 100 Hz by the materials testing system and converted to the engineering stress and strain space following testing using the measured sample height and diameter.

#### 2.1.3 Hyperelastic fitting of fibrin

In ABAQUS (Dassault Systèmes, Vélizy-Villacoublay, France), stress-strain data for confined compression tests (data up to 5.0% strain) were paired with unconfined compression tests (data up to 33% strain) and imported as uniaxial compression and volumetric compression data, respectively. The data were then fit to polynomial (first order to third order), Ogden (first order to third order), reduced polynomial (first order to third order), and Van der Waals compressible hyperelastic models. The most appropriate model for finite element implementation was selected based on model stability and fit to experimental data. The averaged material models (elasticity and compressibility independently) were compared to the experimental data resampled 0.1% strain increments with lack-of-fit F-tests (α = 0.05 for all tests).

### 2.2 Scaffold unit cell model

#### 2.2.1 Geometry and materials

In ABAQUS, a repeating unit of an angle-ply laminate scaffold geometry was generated based on a previously developed scaffold (Figure 2a-b)(8). This angle-ply fiber architeture has demonstrated similar anisotropic mechanical properties to native annulus fibrosus tissue. Polycaprolactone (PCL) fibers were assumed to be linear elastic (E = 265 MPa, v = 0.3) and idealized as straight cylinders. Contact areas at fiber intersections were simplified by generating cylindrical columns through the thickness of the scaffold at each fiber intersection. A hydrogel matrix infill was modelled in the scaffold void spaces (Figure 2c) and prescribed compressible, hyperelastic material properties based on the described characterization of fibrin. The fiber and matrix geometries were merged, assuming perfect bonding between the two materials. At the center of the hydrogel geometry, a region of interest (ROI) of mesh elements was defined as a repeating unit that encompassed all possible positions of the hydrogel matrix relative to the fiber scaffold.

**Figure 2.**
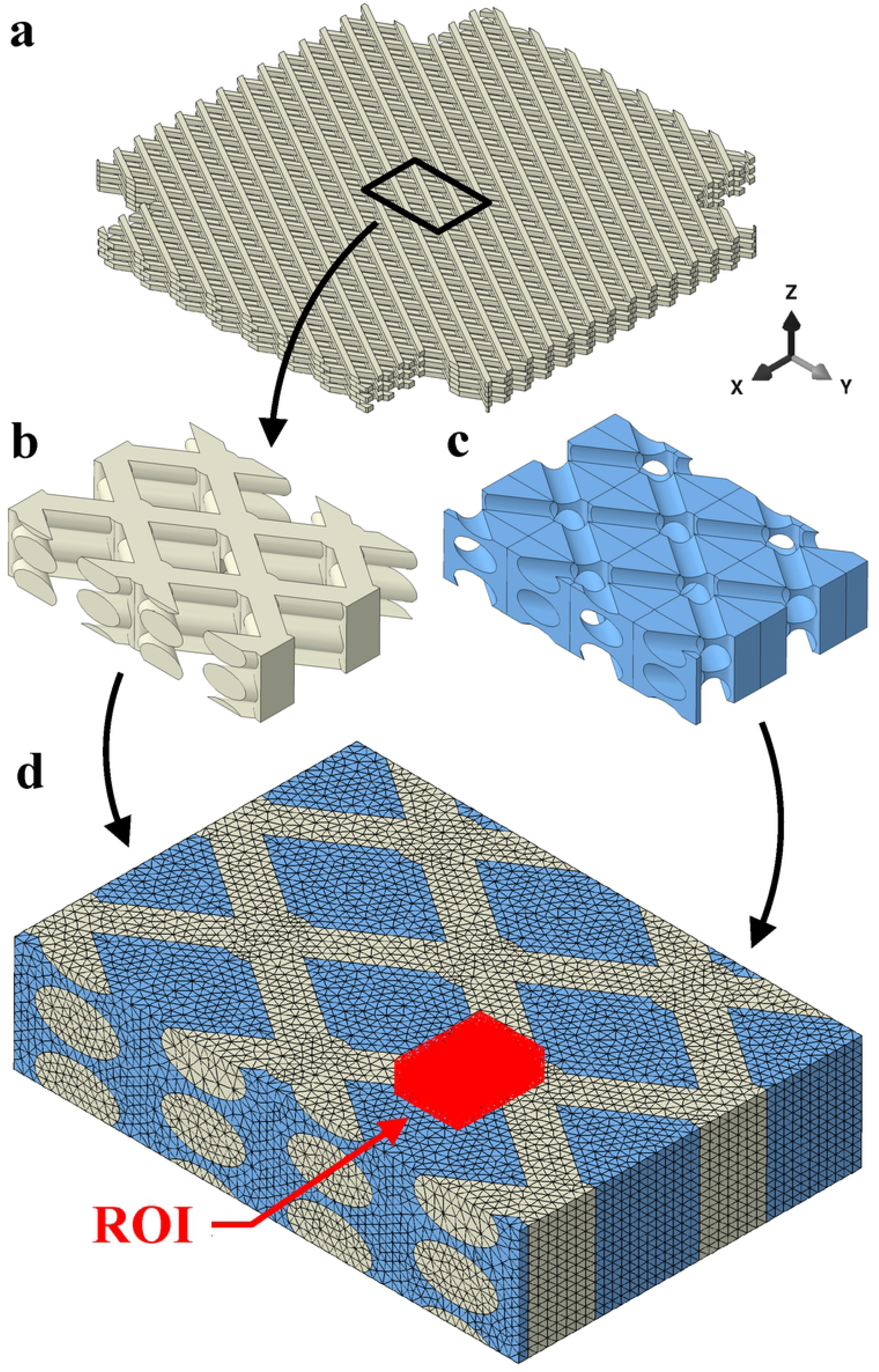
Scaffold model for the base geometry showing (a) the previously validated angle-ply fiber scaffold (8), (b) the double unit cell of the fiber scaffold, (c) the hydrogel infill for the double unit cell, and (d) the final model with mesh and ROI for CME evaluation. The x-, y-, and z-directions represent the axial, circumferential, and radial directions of the IVD, respectively.

This particular model was based on a TE strategy for repair of the annulus fibrosus. Accordingly, to mimic the dominant *in vivo* loads experienced by the posterolateral annulus fibrosus, tensile strains were applied to the model in the x-direction (axial) and y-direction (circumferential) (10,11). For all model validation, 5.0% equibiaxial strains were generated by prescribing displacements on the positive axial and circumferential faces. All ABAQUS jobs were computed on a Linux operating system (CentOS Linux 7 (Core)) with 60 processor cores (Xeon E5 2683 v4, Intel Corporation, Santa Clara, CA) and 128 GB RAM.

### 2.3 Model validation

#### 2.3.1 Unit cell size

The influence of the idealized boundary conditions at the circumferential and axial faces (i.e., uniform normal displacement) on the ROI mechanics was investigated. The double (2-by-2) unit cell was compared to 1-by-1, 1.5-by-1.5, and 3-by-3 unit cells, representing both increases and reductions of the influence of boundary conditions on the ROI mechanics. Specifically, the ROI strain energy (SE) and total unit cell SE were used to measure the ROI mechanics and the central processing unit (CPU) time was used to measure the trade-off in solution time. Preliminary data indicated that a perfectly centered ROI produced unstable boundary conditions because the boundary coincided with small geometric features. This instability was also exacerbated when architectural parameters were changed. Accordingly, the ROI was offset in the positive y-direction by 1/8 of a unit cell to provide more stable boundary geometries whilst minimizing the boundary influence on the ROI mechanics.

#### 2.3.2 Mesh refinement

Validation of the unit cell size was conducted prior to mesh refinement because preliminary data indicated that mesh convergence was dependent on the unit cell size. All unit cell validation was performed with a uniform quadratic tetrahedral (C3D10) mesh with 40 μm seed size, which was subsequently validated. The selected unit cell size was then prescribed a series of mesh sizes to evaluate convergence of the mesh and ROI mechanics. Uniform meshes of 40 μm, 35 μm, and 30 μm seed size were considered. For more coarse meshes, a 40 μm seed size was maintained in the ROI to ensure cell-sized volumes were maintained for CME evaluation. The remaining unit cell was seeded with increasing size from 40 μm to 250 μm. For all meshes, the element growth rate within the ROI was set to 1.0 and the element growth rate outside of the ROI was set to 5%.

#### 2.3.3 Boundary conditions

To further understand the influence of unit cell boundary conditions on the mechanics within the ROI, the base model was tested with numerous boundary conditions. In the radial (out-of-plane) direction, the following conditions were modified: (1) constraining all nodes on the positive and negative radial faces, (2) constraining all nodes on the positive radial face, (3) constraining all nodes on the negative radial face, and (4) changing the bilayer count (N) to N = 1, N = 3 and N = 4 to create scaffolds of different thicknessess. In the axial and circumferential directions, the following conditions were modified: (5) removing constraints from the hydrogel on all four in-plane faces, (6) removing constraints from the hydrogel on both axial direction faces, (7) removing constraints from the hydrogel on both circumferential direction faces.

## 3 Results and Discussion

### 3.1 Hyperelastic characterization of fibrin hydrogel

The only hyperelastic models with extensive elastic and volumetric stability were the first-order reduced polynomial (neo-Hookean; all data fits were stable) and second-order reduced polynomial (the elastic data from one sample was unstable and omitted, resulting in n = 6 for this model). Of these two models, both exhibited no statistically significant evidence for lack of fit to the volumetric data (p = 0.99 for both). Similarly, for the elastic data, there was no statistically significant evidence for lack of fit of the second-order model to the experimental data (p = 0.99). However, the first-order model had statistically significant evidence of lack of fit to the experimental data (p < 0.001). Based on the stability and accuracy measures, the second-order reduced polynomial model was selected for subsequent analyses ((1; Figure 3).

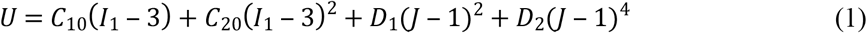

where U is the strain energy potential, C_10_ and C_20_ are the fitted material elasticity constants, I_1_ is the first invariant of the strain tensor, D_1_ and D_2_ are the fitted material compressibility constants, and J is the Jacobian determinant of the deformation gradient tensor. Average fitted coefficients for the second-order reduced polynomial model were obtained by averaging the coefficients from all samples (Table 1).

**Figure 3.**
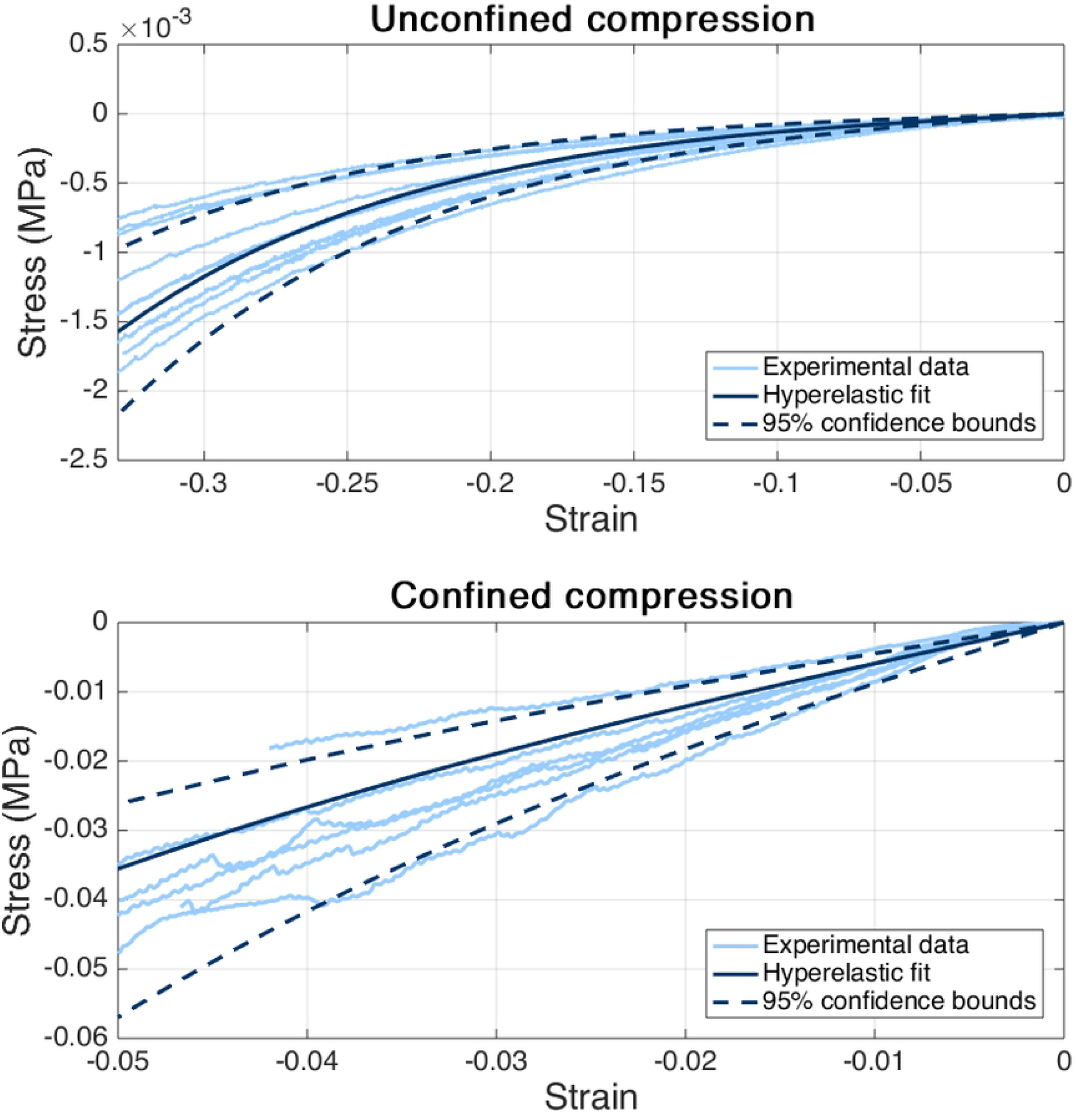
Unconfined and confined compression experimental data with hyperelastic model fits (second-order reduced polynomial). The 95% confidence bounds of the model coefficients (Table 1) are shown (dashed lines). One confined compression sample failed prior to 5% strain, however, was retained in the fitted data set.

**Table 1.** Summary of second-order fitted coefficients for fibrin hydrogel.

| | $C_{10}$ (MPa) | $C_{20}$ (MPa) | $D_1$ (MPa) | $D_2$ (MPa) |
| --- | --- | --- | --- | --- |
| Average | $1.72 \times 10^{-4}$ | $3.83 \times 10^{-4}$ | 3.41 | $8.06 \times 10^{-2}$ |
| Standard deviation | $9.64 \times 10^{-5}$ | $1.76 \times 10^{-4}$ | 1.05 | $4.15 \times 10^{-2}$ |
| Upper 95% confidence bound | $2.46 \times 10^{-4}$ | $5.19 \times 10^{-4}$ | 4.51 | $1.24 \times 10^{-1}$ |
| Lower 95% confidence bound | $9.79 \times 10^{-5}$ | $2.48 \times 10^{-4}$ | 2.30 | $3.71 \times 10^{-2}$ |

### 3.2 Model validation

The results of unit cell validation demonstrated that cell sizes greater than 1.5-by-1.5 had ROI strain energy within 0.01% of the 3 by 3 unit cell (Figure 4). The ROI strain energies of the 1-by-1 unit cells with centered and offset ROI were 38% and 65% greater, respectively, than the 3-by-3 unit cell. Due to the observed influence of geometric boundary conditions on model stability in preliminary studies, the 2-by-2 unit cell was selected as the most suitable size for this study. All three of the reduced boundary conditions on the in-plane (x- and y-direction) faces resulted in changes to the ROI strain energy of 0.5% or less.

**Figure 4.**
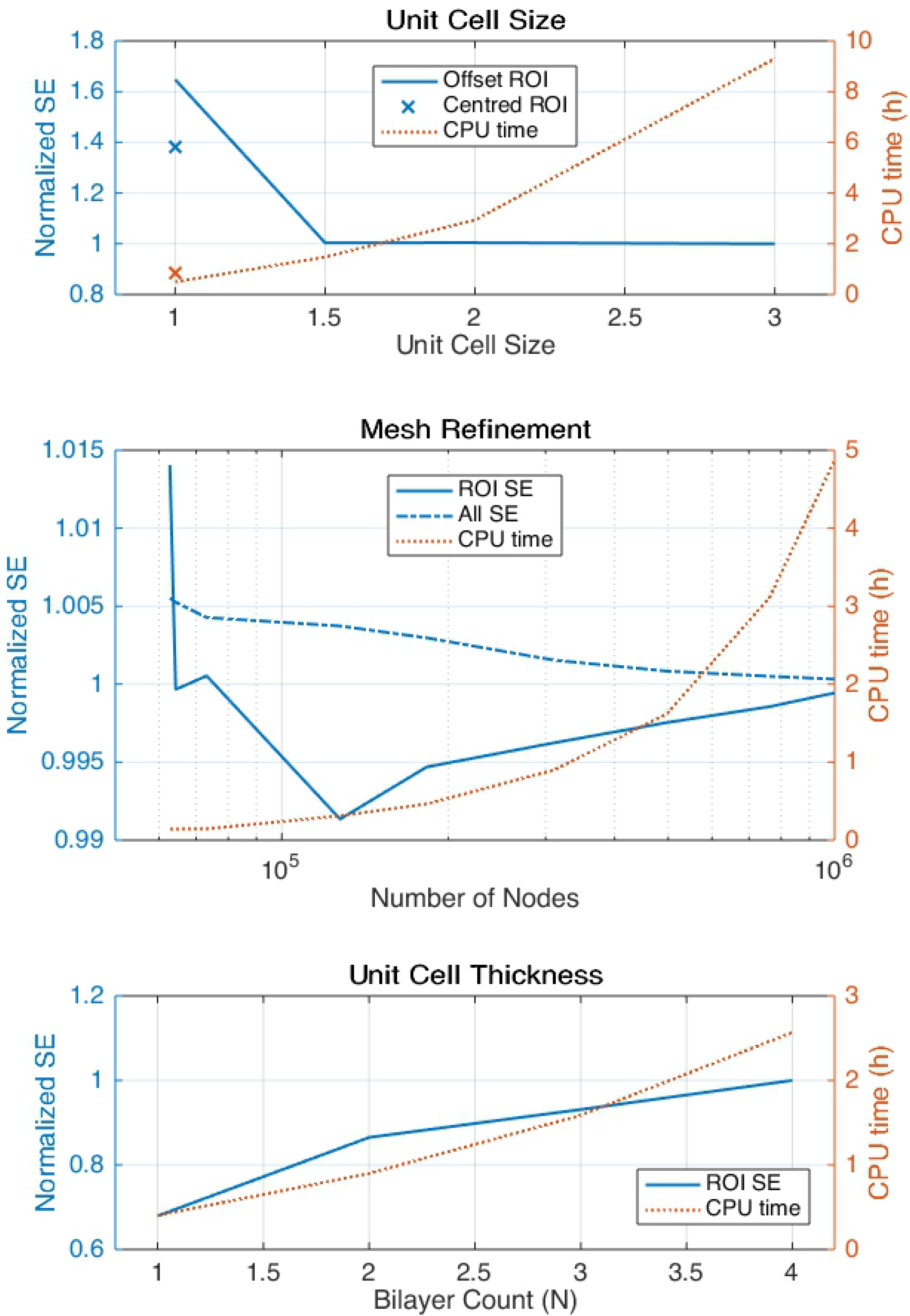
Validation of unit cell size, mesh refinement, and unit cell thickness for the scaffold CME model. For each validation, the convergence of strain energy was weighed against CPU time. In the unit cell validation, a centered ROI was shown to be similar to an ROI offset by 1/8 of a unit cell in order to improve boundary conditions.

The mesh refinement yielded total strain energies within 1% of the 30 μm mesh for all considered meshes. Similarly, the ROI strain energy was within 1% of the 30 μm mesh for all meshes and exhibited apparent convergence for meshes finer than 100 μm. Therefore, the uniform 40 μm mesh was deemed acceptable as compared to finer uniform meshes and the 40 μm ROI mesh was also accepted for all coarser meshes. To select the most appropriate mesh with a 40 μm ROI size, the ROI strain energy and computational time were considered. The 60 μm mesh (0.9 hours CPU time) had ROI strain energy within 0.5% of the uniform 30 μm mesh (10.6 hours CPU time). Therefore, the mesh with 60 μm general size and 40 μm ROI size was selected for all subsequent analyses.

The ROI SE demonstrated a monotonic increase as a function of bilayer count; the one-, two-, and three-bilayer scaffolds yielded SE of 68%, 87%, and 93%, respectively, of the four bilayer scaffold SE. Based on these results, the two bilayer (N = 2) was selected for all subsequent model validation. The fully constrained boundary condition on both out-of-plane (z-direction) faces failed to converge. When considered independently, the fully constrained boundary condition on the top and bottom faces resulted in increases to the ROI strain energy of 179% and 196%, respectively. All of the modified boundary conditions in the out-of-plane direction (z-direction) yielded changes in the total model strain energy of 0.03% or less.

Based on the results of all considered boundary conditions, the out-of-plane (z-direction) boundary condition will likely have the greatest influence on the ROI CME. Careful consideration should be given to the context of these imposed constraints, for example, how these constraints relate to TE scaffolds both *in vitro* and *in vivo*. These constraints are also idealized as uniform planar deformation, which may not reflect the true boundary conditions in a physical scaffold. The scaffold boundary conditions were constrained in this manner to balance ROI convergence, numerical stability, and computational time. Periodic BCs are an alternative approach to mitigate the influence of the boundaries on the ROI (12). However, periodic boundary conditions add considerable computational burden, and, because ROI convergence was demonstrated for the BCs in the current study, periodic boundary conditions may not impart an improvement to the accuracy of results.

Overall, this study demonstrated the convergence of the ROI mechanics in a unit cell model as a function of the unit cell size and mesh refinement. The uniform mesh in the ROI provides a basis to evaluate the CME in all possible cell-sized volumes within the scaffold. This model will subsequently be used to investigate the relative influence of the scaffold loading, materials, and architecture on the CME. Ultimately, this model will leveraged to predict cell local cell fate and guide the design of TE scaffolds for enhanced regeneration.

## 4 Acknowledgements

The authors would like to acknowledge the Preclinical Surgical Research Laboratory at Colorado State University for their assistance in collection of ovine blood for this study. The authors also acknowledge the funding support of the J.R. Templin Trust.

